# An Enhanced Split Intein-Mediated Ligation (SIML) Platform for Rapid Discovery and Functional Screening of Circular Bacteriocins

**DOI:** 10.64898/2026.05.21.726877

**Authors:** Ester Sevillano, Mohamed El Bakkoury, Irene Lafuente, Nuria Peña, Cleopatra Collado, Luis M. Cintas, Estefanía Muñoz-Atienza, Philippe Gabant, Pablo E. Hernández, Juan Borrero

**Author notes:** **Correspondence: Corresponding Authors Estefanía Muñoz-Atienza** – Departamento de Nutrición y Ciencia de los Alimentos (NUTRYCIAL), Sección Departamental de Nutrición y Ciencia de los Alimentos (SD-NUTRYCIAL), Facultad de Veterinaria, Universidad Complutense de Madrid (UCM), Avenida Puerta de Hierro, s/n, 28040 Madrid, Spain, **Juan Borrero** – Departamento de Nutrición y Ciencia de los Alimentos (NUTRYCIAL), Sección Departamental de Nutrición y Ciencia de los Alimentos (SD-NUTRYCIAL), Facultad de Veterinaria, Universidad Complutense de Madrid (UCM), Avenida Puerta de Hierro, s/n, 28040 Madrid, Spain.

## Abstract

Bacteriocins are ribosomally synthesized antimicrobial peptides with promising applications in biotechnology, particularly in food preservation and animal and human health. Circular bacteriocins are especially attractive due to their head-to-tail cyclized structure, which confers enhanced stability and antimicrobial potency relative to linear peptides. Here, we report an *in vitro* cell-free protein synthesis system coupled with an enhanced Split Intein-Mediated Ligation platform (IV-CFPS/SIML) for the efficient synthesis of circular bacteriocins through systematic evaluation of cyclization sites and alternative split inteins. Using enterocin AS-48 as a model, we systematically evaluated multiple serine-based cyclization sites in combination with three split inteins, NpuDnaE, Gp41-1, and SspGyrB, to identify configurations supporting efficient splicing and high antimicrobial activity. Gp41-1 emerged as the most effective intein and was subsequently applied to the production of garvicin ML, amylocyclicin, and 27 naturally occurring sequence variants, demonstrating that cyclization site selection, intein identity, and minor sequence variations strongly influence antimicrobial potency and target range. Finally, SIML expression cassettes encoded in pUC-derived vectors enabled *in vivo* production and functional expression of selected circular bacteriocins in recombinant *Escherichia coli*. Collectively, these results establish SIML as a versatile platform for *in vitro* and *in vivo* production, screening, and functional characterization of known and putative circular bacteriocins.

## 1. Introduction

The rapid emergence of antibiotic-resistant pathogens represents a major global health threat, underscoring the urgent need for novel antimicrobial agents [1,2]. In this context, bacteriocins, ribosomally synthesized antimicrobial peptides produced by bacteria, have attracted considerable attention as promising alternatives to conventional antibiotics due to their high specificity, potent antimicrobial activity, low toxicity toward eukaryotic cells, and effectiveness against antibiotic-resistant bacteria [3]. Bacteriocins are commonly classified based on their structure, mode of action, and the presence or absence of post-translational modifications. Class I bacteriocins, which undergo post-translational modifications, include several subclasses, including the head-to-tail cyclized bacteriocins known as circular bacteriocins. These peptides undergo a unique N-to-C terminal ligation that generates a compact globular structure composed of multiple α-helical motif surrounding a hydrophobic core, conferring exceptional resistance to heat, extreme pH conditions, and proteolytic degradation [4–6].

Head-to-tail cyclized bacteriocins are encoded in biosynthetic gene clusters comprising genes for the precursor peptide, transporter(s), a SpoIIM (DUF95) membrane protein, an immunity protein, and additional hydrophobic proteins [6,7]. They are synthesized as linear precursors with an N-terminal leader that is proteolitically removed, followed by head-to-tail cyclization of the core peptide prior to secretion. Despite this general framework, the molecular mechanisms governing leader peptide cleavage, peptide cyclization, and secretion remain incompletely understood [6–9]. Circular bacteriocins primarily exert antimicrobial activity through direct, receptor-independent interaction with the target cell membrane, leading to dissipation of membrane potential, ion leakage, and ultimately cell death. However, some circular bacteriocins may also display a dual, concentration-dependent mode of action involving receptor-mediated recognition at lower concentrations [6,7].

New circular bacteriocins have traditionally been identified through activity-guided purification from native producer strains or by heterologous expression, approaches that require extensive gene cloning and peptide purification. These methods are labor-intensive, time-consuming, and often rely on the co-expression of additional genes involved in bacteriocin maturation and secretion [6,7,10–12]. To overcome these limitations, we explored the use of split inteins, which have long been applied to drive peptide and protein cyclization [13]. Building on this strategy, our group previously demonstrated that the split NpuDnaE intein from *Nostoc punctiforme* enables rapid and reliable production of circular bacteriocins both *in vitro* and *in vivo* using *E. coli* as heterologous host [14,15] (Fig. 1). Notably, the SIML system circumvents the need for additional biosynthetic genes and enables the efficient synthesis of biologically active circular bacteriocins. The use of an IV-CFPS system coupled with SIML (IV-CFPS/SIML), in combination with the split NpuDnaE intein, has facilitated the production of nearly 20 previously described or putative circular bacteriocins, underscoring the robustness, versatility, and broad applicability of this platform [14].

**Fig. 1.**
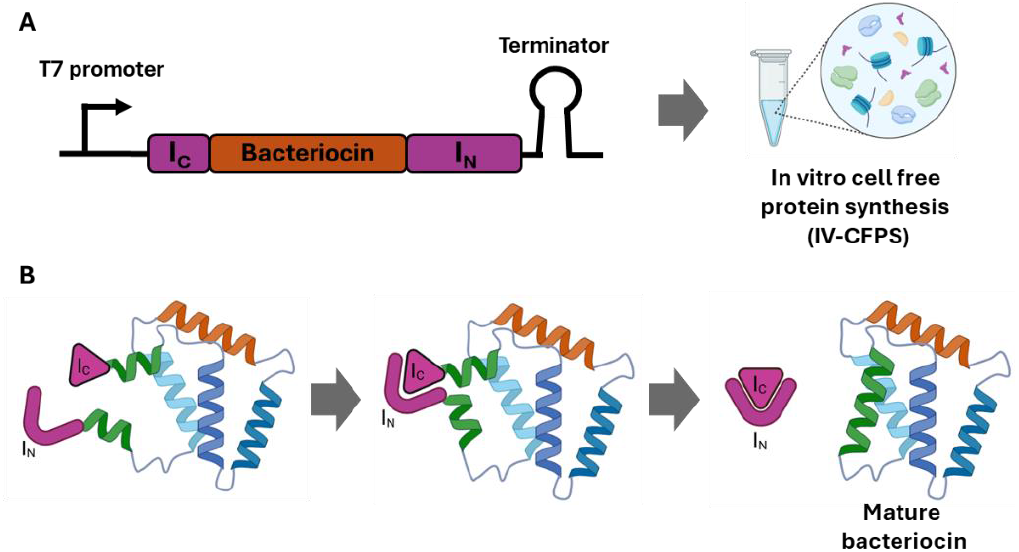
Schematic overview of the SIML workflow for *in vitro* production of circular bacteriocins using an IV-CFPS/SIML platform. (A) A synthetic gene encoding the mature circular bacteriocin is cloned between the C-terminal (I_C_) and N-terminal (I_N_) halves of a split intein. The synthetic gene construct is expressed using an IV-CFPS system, yielding a linear fusion precursor containing both intein fragments. (B) Upon folding, the split intein halves associate to form an active intein domain that catalyzes peptide cyclization. Following intein fragment association, trans-splicing proceeds through the canonical (thio)esterification and rearrangement steps, resulting in intein excision and formation of a head-to-tail cyclized bacteriocin.

Split inteins are naturally occurring protein elements that mediate protein splicing, a self-catalyzed process in which the intein excises itself while covalently ligating the flanking peptide segments through formation of a new peptide bond. In addition to NpuDnaE, several other split inteins, including Gp41-1 and SspGyrB, have been characterized for their high splicing efficiency and minimal sequence requirements, making them powerful tools for peptide and protein cyclization [16–18]. The Gp41-1 intein, identified within a phage-related *gp41* gene from environmental metagenomic sequences, exhibits exceptionally fast splicing kinetics and remarkable robustness across diverse conditions [19,20]. Similarly, the SspGyrB intein from *Synechocystis* sp. PCC 6803 has been widely employed in protein engineering owing to its high efficiency and broad tolerance to sequence variation [17,21–23]. These highly active inteins therefore represent attractive alternatives to NpuDnaE for site-specific peptide cyclization, providing an opportunity to systematically evaluate how cyclization-site selection and intein identity influence circular bacteriocin production and activity. Importantly, split intein performance is strongly influenced by the identity of the extein residues flanking the splicing junction, particularly the +1 C-extein residue, which must be a nucleophilic amino acid such as Cys, Ser, or Thr [24]. Because extein composition directly affects splicing efficiency and product yield, careful selection of both the cyclization site and split intein is essential for successful circular peptide engineering [25–28].

In this work, we systematically investigated how cyclization site-selection and intein identity influence the production and antimicrobial activity of circular bacteriocins using the IV-CFPS/SIML platform. Using enterocin AS-48 (EntAS-48) as a model, multiple serine-based cyclization sites were evaluated in combination with three split inteins (NpuDnaE, Gp41-1, and SspGyrB) to identify configurations that support efficient splicing and high antimicrobial activity. The optimized IV-CFPS/SIML platform was subsequently extended to the production of garvicin ML (GarML), amylocyclicin (AmCyc), and 27 sequence variants to determine how cyclization-site selection, intein choice, and minor sequence variations influence antimicrobial activity and target range. Finally, SIML expression cassettes encoded into pUC-derived protein expression vectors were evaluated for the *in vivo* production, cyclization, and functional expression of selected circular bacteriocins in recombinant *Escherichia coli*.

## 2. Methods

### 2.1. Construction of SIML Cassettes for IV-CFPS Expression

SIML expression cassettes were generated by overlap-extension PCR using a three-fragment assembly strategy (Fig. S1). Fragment A consisted of a synthetic DNA sequence containing the T7 promoter followed by the C-terminal fragment of the corresponding split intein (I_C_). Fragment B comprised a synthetic DNA sequence encoding the mature circular bacteriocin sequence, starting at the selected serine residue, and flanked by 5′ homology to I_C_ and 3′ homology to the N-terminal intein fragment (I_N_). Fragment C consisted of a synthetic DNA sequence containing the I_N_ intein fragment followed by a transcription terminator. All sequences were reverse-translated using *E. coli* codon usage (GeneArt Gene Synthesis tool) and synthesized by GeneArt (ThermoFisher Scientific). Fragments A, B, and C were assembled by overlap-extension PCR to produce a complete T7–I_C_–bacteriocin–I_N_–terminator expression cassette. Final PCR products were purified using the Nucleospin Gel and PCR Clean-up kit (Macherey-Nagel) and used directly as DNA templates for IV-CFPS reactions.

For the design of EntAS-48 constructs with alternative cyclization sites, four internal serines (S30, S37, S41, and S50) were individually selected as the +1 nucleophile for split intein-mediated circularization (Table S1). To evaluate alternative split inteins, equivalent SIML-derived expression cassettes were generated in which the mature peptide sequence was flanked by either Gp41-1 or SspGyrB split intein instead of NpuDnaE. For each intein, four constructs were produced by independently assigning S30, S37, S41, or S50 as the +1 cyclization residue (Table S1). For GarML and AmCyc, SIML expression cassettes were generated using the split Gp41-1 intein. Multiple constructs were designed by individually selecting internal serine residues as the +1 cyclization site (S19, S29, and S32 for GarML; S3, S8, S23, and S30 for AmCyc) (Table S1). Sequence variants of EntAS-48, GarML, and AmCyc were identified by BLASTp analysis [29] and selected based on overall sequence identity and the presence of a conserved serine residue at the cyclization position of the corresponding reference bacteriocin. This approach yielded nine EntAS-48 variants (containing S41 as the +1 residue), four GarML variants (containing S29 at position +1), and eleven AmCyc variants (containing S23 at position +1). For each variant, SIML expression cassettes were generated by synthesizing Fragment B starting at the conserved serine residue and assembling it with Fragments A and C as described above. In total, 27 sequence variants were constructed and designated AS48_1–AS48_10 (EntAS-48), GML_1–GML_5 (GarML variants), and ACC_1–ACC_12 (AmCyc variants), as summarized in Table S1.

### 2.2. *In Vitro* Production of Circular Bacteriocins (IV-CFPS/SIML)

PCR-assembled SIML expression cassettes (T7–I_C_–bacteriocin–I_N_–terminator), generated as described above, were used as purified DNA templates for IV-CFPS. Reactions were performed using the PURExpress® *In Vitro* Protein Synthesis Kit (New England Biolabs, Ipswich, MA, USA) according to the manufacturer instructions, with minor modifications. Each 25 µL IV-CFPS reaction contained PURExpress Solutions A and B, nuclease-free water, and purified expression cassette DNA at a final concentration of 10 ng µL^-1^. Reactions were incubated at 37 °C for 2 h and subsequently held at room temperature (20–22 °C) overnight to allow completion of split intein–mediated trans-splicing and head-to-tail peptide cyclization. Unless otherwise stated, IV-CFPS/SIML reaction mixtures were used directly for downstream antimicrobial activity assays.

### 2.3. Evaluation of Antimicrobial Activity of IV-CFPS/SIML Products

The antimicrobial activity of EntAS-48 circularized at positions S30, S37, S41, and S50 using the split Npu DnaE intein was evaluated by the spot-on-agar test against *P. damnosus* CECT 4797, as previously described [30]. Two-fold serial dilutions of each IV-CFPS/SIML reaction mixture were prepared, and 5 μL aliquots of each dilution were spotted onto de Man, Rogosa and Sharpe (MRS) agar plates (1.5% w/v) overlaid with soft agar (0.8% w/v) inoculated with approximately 10^5^ CFU mL^-1^ of the indicator strain. Plates were incubated at 30 °C for 24 h, after which inhibition zones were recorded. The spot-on-agar assay was also used to assess the antimicrobial activity of GarML and AmCyc constructs produced via IV-CFPS/SIML using different serine cyclization sites and the Gp41-1 intein. In addition, spot-on-agar assays were performed to evaluate the antimicrobial activity of the sequence variants of EntAS-48 (AS48_1-AS48_10), GarML (GML_1–GML_5), and AmCyc (ACC_1–ACC_12) against a broader panel of Gram-positive and Gram-negative indicator strains (Table S2).

The antimicrobial activity of samples was also quantified using a microtiter plate assay (MPA) against *P. damnosus* CECT 4797, as previously described [31]. IV-CFPS/SIML reaction mixtures containing EntAS-48 cyclized at S30, S37, S41, or S50 using NpuDnaE, Gp41-1, or SspGyrB were serially diluted two-fold in 96-well microplates. Bacterial growth was monitored spectrophotometrically at 620 nm using a FLUOstar OPTIMA plate reader (BMGLabtech, Ortenberg, Germany). One bacteriocin unit (BU) was defined as the reciprocal of the highest dilution producing 50% inhibition growth of the indicator strain [32]. Results are reported as the mean ± standard error of three independent experiments.

### 2.4. *In Vivo* Production, Purification, and Proteomic Characterization of Circular Bacteriocins

To evaluate *in vivo* production of SIML-derived circular bacteriocins, synthetic gene fragments encoding bacteriocins EntAS-48 (AS48_1; S41) and AmCyc variant 3 (ACC_3; S23), flanked by the C-terminal (I_C_) and N-terminal (I_N_) and fragments of the Gp41-1 intein, were cloned into pUC-derived protein expression vectors. These vectors contain a T7 promoter, start codon (ATG), stop codon (TAA), and T7 transcription terminator, yielding the plasmids pCirc-Gp41-AS48_1 (S41) and pCirc-Gp41-ACC_3 (S23), synthesized by GeneArt (Thermo Fisher Scientific). Transformation of competent *E. coli* cells with these plasmids generated *E. coli* BL21 (DE3) (pCirc-Gp41-AS48_1 (S41)) and *E. coli* BL21 (DE3) (pCirc-Gp41-ACC_3 (S23) strains for intracellular production of the bacteriocins AS48_1 and ACC_3, respectively.

The expression, purification, and antimicrobial activity assessment of intracellular bacteriocins were performed as previously described [14,15], with minor modifications. Briefly, 1 L of LB broth supplemented with kanamycin (50 µg mL^-1^) was inoculated with an overnight culture of the corresponding recombinant strain and incubated at 37 °C with shaking at 250 rpm until an OD_620_ of approximately 0.1 was reached. Cultures were grown to an OD_620_ of ∼0.4, induced with 0.5 mM IPTG, and incubated for an additional 3 h. Cells were harvested by centrifugation (8,000 × g, 15 min, 4 °C), resuspended in 40 mL of ice-cold 20 mM phosphate buffer (pH 6.0) containing 1 M NaCl, and lysed by sonication using a Branson 450 Digital Sonifier (six cycles of 10 s at 45% amplitude, with 1 min cooling on ice between pulses). Insoluble material was removed by centrifugation (8,000 × g, 15 min, 4 °C), and the clarified soluble fraction was filtered through a 0.45 µm membrane (Sartorius).

Bacteriocins AS48_1 and ACC_3 were purified from the cellular-soluble fraction by hydrophobic interaction chromatography (HIC) on Octyl-Sepharose CL-4B, followed by reverse-phase fast protein liquid chromatography (RP-FPLC) using an ÄKTA Pure system (GE Healthcare Life Sciences), as previously described [15]. Samples were applied to a Resource RPC 3 mL column (Cytiva) and eluted with a linear gradient of 0–100% isopropanol containing 0.1% (v/v) trifluoroacetic acid (TFA). Elution was monitored by UV absorbance at 254 nm, and antimicrobial activity of collected fractions was quantified using a microtiter plate assay against *P. damnosus* CECT 4749 [32]. Fractions exhibiting the highest antimicrobial activity were analyzed by MALDI–TOF mass spectrometry for molecular mass determination at the Proteomics Facility (CAI Técnicas Biológicas, Complutense University of Madrid, Spain), following established procedures [32].

## 3. Results and discussion

The global rise of antibiotic-resistant pathogens has intensified the search for innovative antimicrobial strategies, among which circular bacteriocins are particularly attractive due to their exceptional stability and potent activity. To overcome limitations in their rapid screening and production, we developed a synthetic biology platform based on SIML, which enables head-to-tail cyclization of bacteriocins independently of native biosynthetic machinery. Notably, SIML permits the production of mature circular bacteriocins without the need for additional gene cluster-encoded proteins involved in processing, transport, regulation, or immunity [14,15]. Here, we systematically investigate how cyclization-site selection and split intein identity influence the antimicrobial activity of SIML-derived bacteriocins.

### 3.1. Influence of Circularization Site on Enterocin AS-48 Antimicrobial Activity

Because intein performance is strongly influenced by the identity of the extein residues flanking the splicing junction, we first examined the biochemical constraints governing the +1 position. Intein-mediated splicing is most efficient when a nucleophilic residue occupies the +1 extein position (typically Cys, Ser, or Thr) whose steric and electronic properties strongly influence the efficiency of the trans-(thio)esterification reaction that drives peptide bond formation [24]. Among these residues, cysteine is generally the most efficient +1 nucleophile due to the higher reactivity of the thioester intermediate. However, because circular bacteriocins typically lack cysteine residues, serine represents the most favourable alternative. Serine is often preferred over threonine because its smaller side chain imposes fewer steric constraints within the intein active site, facilitating more efficient catalysis. This preference is consistent with surveys of natural inteins, where Ser+1 residues occur more than twice as frequently as Thr+1 (29% vs 12%) [33].

In previous SIML-based studies of circular bacteriocins, the serine residues selected as +1 nucleophiles were chosen without systematic optimization [14,15]. Although these studies demonstrated that SIML enables the production of diverse circular bacteriocins, they did not assess the extent to which alternative serine positions at the cyclization junction contribute to functional variability. Here, we address this gap by systematically evaluating how the position of the serine residues affects intein performance and antimicrobial potency.

As an initial model bacteriocin for this study, we focused on EntAS-48, one of the most extensively characterized circular bacteriocins [34]. The mature 70-amino-acid peptide contains four serine residues (S30, S37, S41, and S50) distributed along its sequence (Fig. 2A), providing a controlled and biologically relevant framework to assess positional effects. Each serine residue was independently elected as the +1 nucleophile in the enhanced IV-CFPS/SIML platform using the split NpuDnaE intein (Fig. 2B), and the antimicrobial activity of the resulting circular bacteriocins was evaluated by spot-on-agar assays against *Pediococcus damnosus* CECT 4797 (Fig. 2C).

**Fig. 2.**
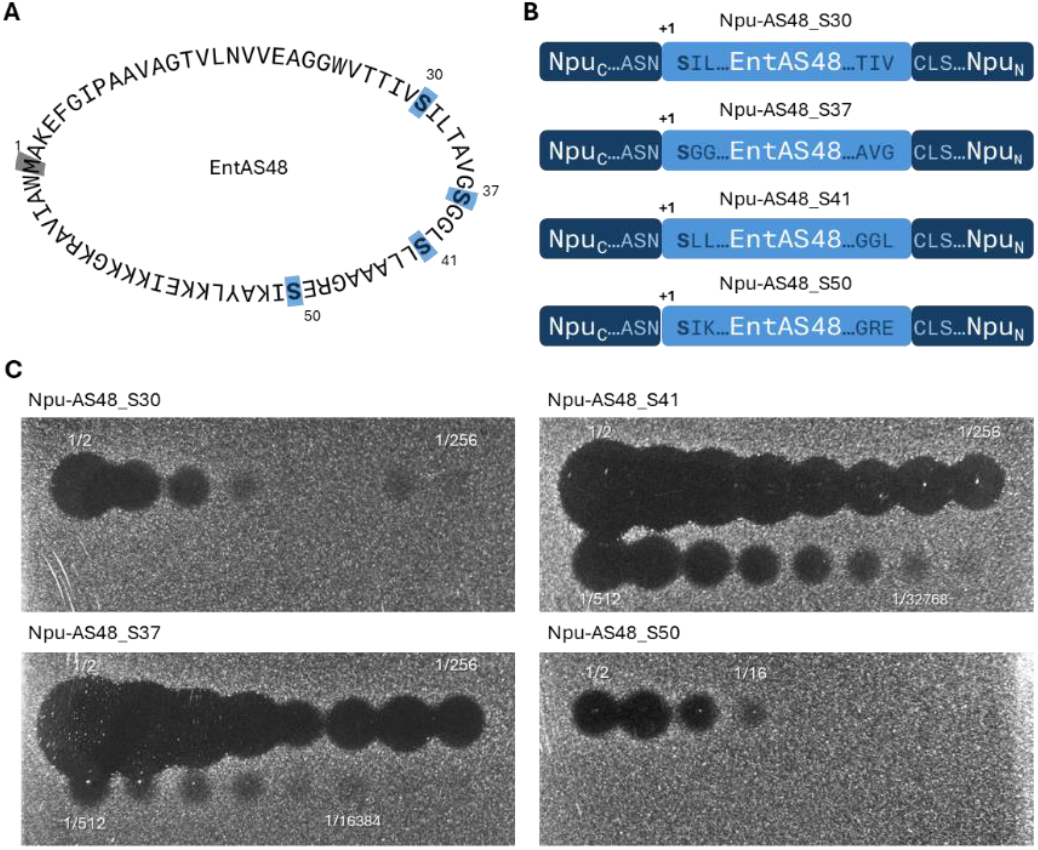
Effect of alternative serine positions on EntAS-48 circularization and antimicrobial activity. (A) Circular representation of EntAS-48 highlighting the four internal serines evaluated as +1 nucleophiles (cyan) and the native cyclization site (grey). (B) Schematic representation of the corresponding SIML expression cassettes constructed using with the split NpuDnaE intein. (C) Spot-on-agar antimicrobial assays of the IV-CFPS/SIML products against *P. damnosus* CECT 4797. Samples were tested using serial two-fold dilutions ranging from 1:2 to 1:65536.

Marked differences were observed among the EntAS-48 circularization-site variants. Npu-EntAS48_S41 exhibited the highest antimicrobial activity, detectable at dilutions up to 1:32,768 of the IV-CFPS/SIML reaction mixture, followed by Npu-EntAS48_S37 (1:16,384), Npu-EntAS48_S30 (1:256), and Npu-EntAS48_S50 (1:16) (Fig. 2C). These results demonstrate that cyclization-site selection profoundly influences functional output, with more than a 4,000-fold difference in activity between the most and least active constructs. Given the sensitivity of intein catalysis to the local extein environment, the S41 junction likely provides a favorable conformational and hydrogen-bonding content that promotes efficient trans-splicing and proper folding of the mature peptide. In contrast, cyclization at S50 may impose steric constraints or unfavorable local topology that limit splicing efficiency, resulting in incomplete ligation and reduced antimicrobial activity. Overall, these findings highlight that even subtle changes at the cyclization junction can markedly affect circularization efficiency and bioactivity., underscoring the importance of rational ligation-site selection in circular bacteriocin engineering.

### 3.2. Influence of Cyclization Site and Split Inteins on Enterocin AS-48 Activity

In previous efforts to develop the SIML platform for circular bacteriocin production [14,15], we employed the split NpuDnaE intein due to its high catalytic efficiency, rapid splicing kinetics, and broad extein tolerance [35,36]. Although native NpuDnaE utilizes cysteine as the +1 nucleophile, it can tolerate serine or threonine at this position, albeit with reduced reaction rates and a stronger dependence on the surrounding extein sequence context [24,36]. Nevertheless, given that NpuDnaE catalysis is intrinsically optimized for cysteine-mediated splicing, we sought to evaluate whether split inteins that naturally employ serine as the +1 extein residue might exhibit improved performance within the SIML framework. Gp41-1 is an ultrafast and highly promiscuous split intein that natively uses serine at the +1 position and is considered among the fastest and most extein-tolerant tolerant inteins characterized to date [18,20]. Similarly, the *Synechocystis* sp. GyrB split intein (SspGyrB) is a high-performance intein with broad sequence tolerance that also relies on serine as the +1 nucleophile [18,37]. Because both Gp41-1 and SspGyrB are intrinsically adapted for serine-driven reactivity, we hypothesized that these inteins would outperform NpuDnaE in catalyzing EntAS-48 cyclization at selected serine ligation sites.

Accordingly, the antimicrobial activity of EntAS-48 cyclized at four serine ligation sites (S30, S37, S41, and S50) was evaluated using the split inteins NpuDnaE, Gp41-1 or SspGyrB in the IV-CFPS/SIML platform (Fig. 3A). The antimicrobial activity of the resulting EntAS-48 peptides varied markedly as a function of both the cyclization site and the intein employed. Among all constructs, Gp41-1-EntAS48_S41 exhibited the highest antimicrobial activity, reaching 1.75 × 10^7^ bacteriocin units (BU), several orders of magnitude greater than any other combination, followed by Npu-EntAS48_S41 (2.18 × 10^6^ BU). Intermediate activity levels were observed for Gp41-1-EntAS48_S37 and Npu-EntAS48_S37, each yield in approximately 4.37 × 10^5^ BU. The lowest antimicrobial activity was observed for SspGyrB-EntAS48_S30 (3.2 × 10^2^ BU). Within the SspGyrB-derived constructs, SspGyrB-EntAS48_S37 displayed the highest activity (4.37 × 10^5^ BU), whereas cyclization at SspGyrB-EntAS48_S41 (1.37 × 10^4^ BU) or SspGyrB-EntAS48_S50 (1.02 × 10^4^ BU) resulted in substantially reduced activity. A Log_10_ reduction of the antimicrobial activity of the EntAS-48 peptide sequences highlights the pronounced dependence of their bioactivity on both the cyclization site and the split intein employed (Fig. 3B).

**Fig. 3.**
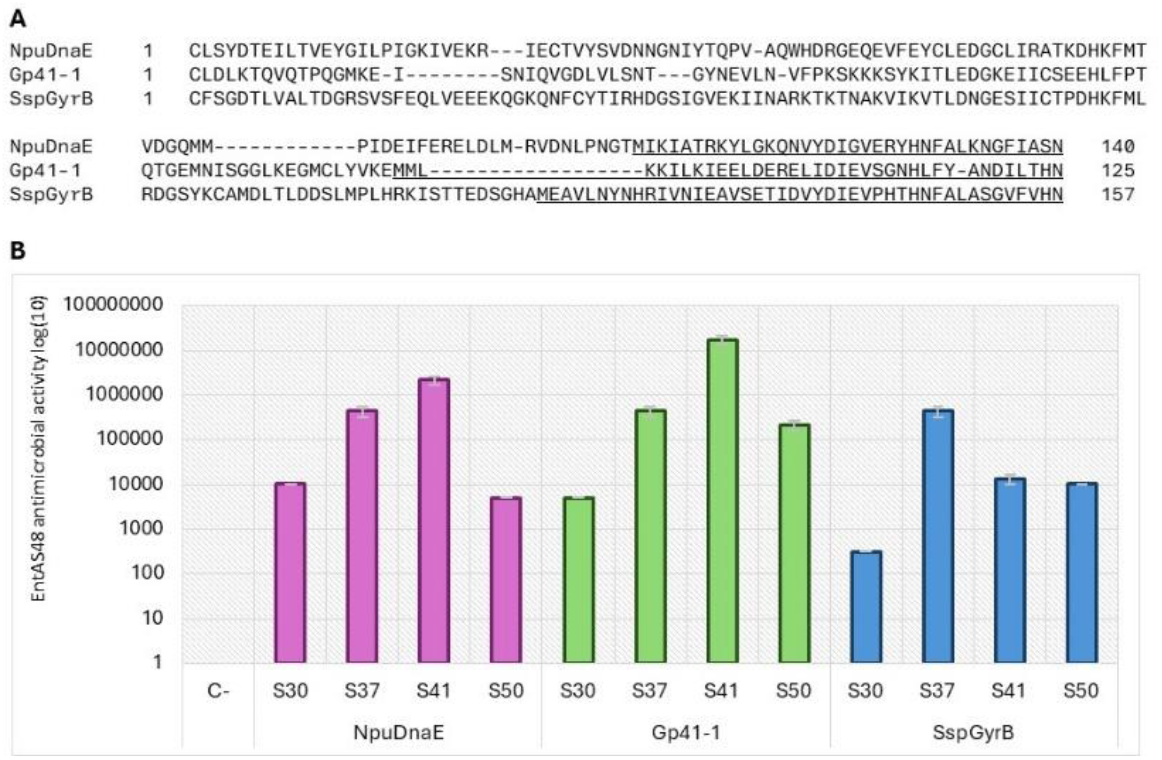
(A) Multiple sequence alignment of the full-length inteins NpuDnaE, Gp41-1, and SspGyrB, with the C-terminal intein fragment (I_C_) underlined. (B) Antimicrobial activity, expressed in bacteriocin units (BU), of EntAS-48 cyclized at four distinct serine residues using three alternative split inteins and the IV-CFPS/SIML platform. EntAS-48 was cyclized at serine positions S30, S37, S41, and S50 using the split inteins NpuDnaE (purple), Gp41-1 (green), or SspGyrB (blue). Antimicrobial activity against *P. damnosus* CECT 4797 is presented on a log_10_ scale; error bars represent the standard error of the mean (n = 3).

These findings demonstrate that intein identity, like cyclization-site selection, is a critical determinant of functional circular bacteriocin production. The exceptional performance of Gp41-1-EntAs48_ S41 is consistent with the ultrafast splicing kinetics and broad extein tolerance of Gp41-1 [18,20], as well as its native reliance on serine as the +1 nucleophile. The alignment between intein catalytic chemistry and the engineered ligation junction likely enhances both splicing efficiency and reaction fidelity. In contrast, the lower performance of SspGyrB-derived constructs suggests that this intein imposes additional structural or sequence constraints that are suboptimal for the compact topology of EntAS-48, despite its robust performance in other protein engineering contexts [18]. Consistent with previous reports, these results highlight the strong dependence of intein activity on local extein conformation and structural context beyond the canonical ™1/+1 residues [17,38]. Overall, these results underscore that intein selection is as critical as cyclization-site selection for achieving high antimicrobial activity in IV-CFPS/SIML systems. They further highlight the necessity of aligning intein catalytic preferences with the sequence and structural features of the target peptide when designing synthetic biology strategies for circular bacteriocin engineering and production.

### 3.3 Influence of Cyclization Site and the Gp41-1 Split Intein on Garvicin ML and Amylocyclicin Activity

The results obtained with EntAS-48 prompted us to investigate whether other circular bacteriocins exhibit a similar sensitivity to cyclization-site selection when produced using the Gp41-1 split intein within the IV-CFPS/SIML platform. Head-to-tail bacteriocins are classified into two subgroups: subgroup I, which is highly cationic with a pI >9, and subgroup II, which is highly hydrophobic with pI <7 [7]. Circular bacteriocins of subgroup I share a conserved saposin-like fold despite substantial sequence diversity [7]. Accordingly, we selected GarML from *Lactococcus garvieae* DCC43 [39], and AmCyc from *Bacillus amyloliquefacien*s FZB42 [40], as structurally related yet sequence-divergent representatives for comparative analysis.

Accordingly, GarML and AmCyc were cyclized at all internal serine residues using the Gp41-1 split intein within the IV-CFPS/SIML platform. GarML was cyclized at serine positions S19, S29, and S32, whereas AmCyc was cyclized at serine positions S3, S8, S23, and S30 (Fig. 4A, B). The antimicrobial activity of the resulting IV-CFPS/SIML products was evaluated against *P. damnosus* CECT 4797 using spot-on-agar assays (Fig. 4C).

**Fig. 4.**
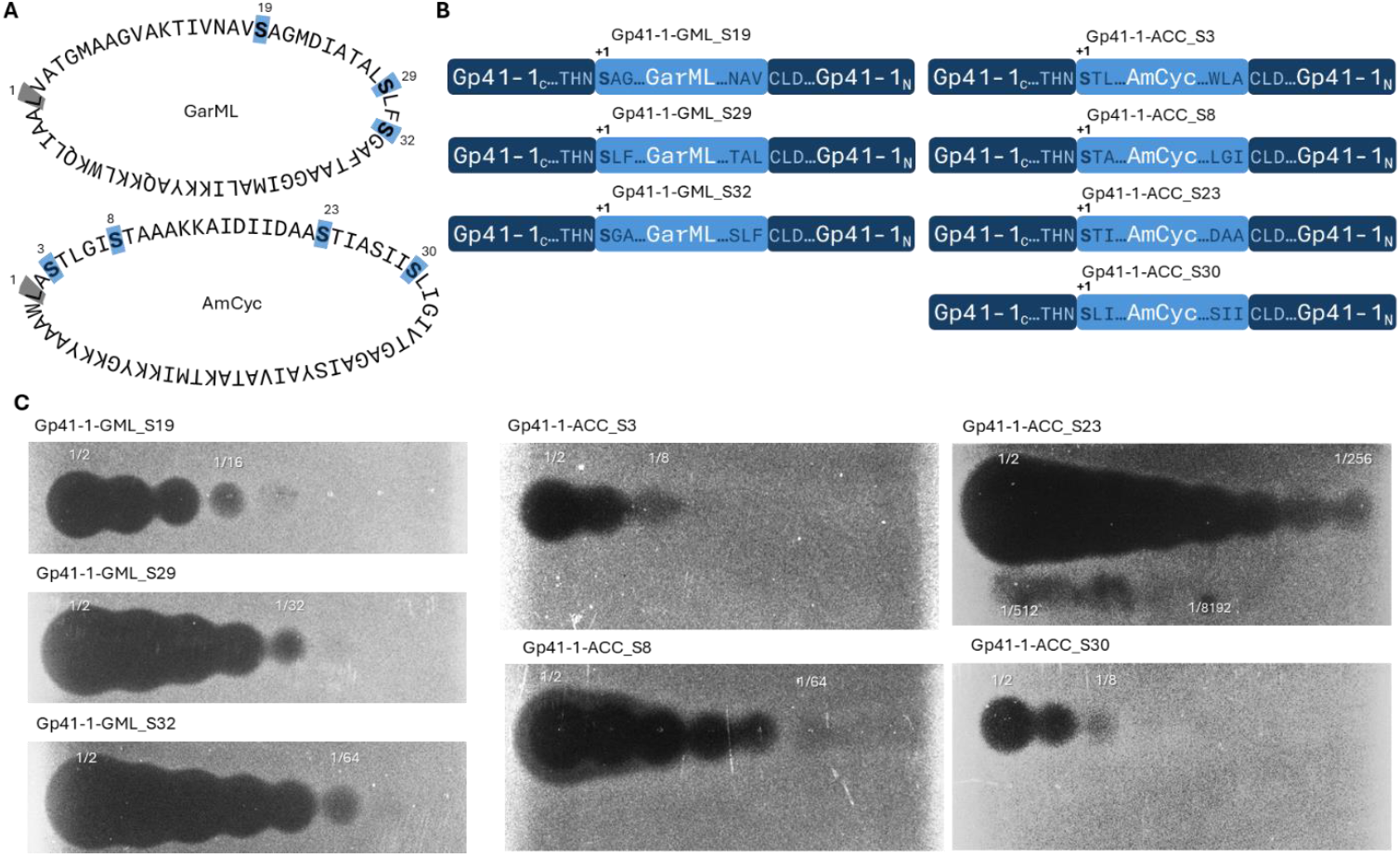
Effect of alternative serine positions on GarML and AmCyc cyclization using the Gp41-1 split intein and the IV-CFPS/SIML platform. (A) Circular representations of GarML and AmCyc highlighting internal serine residues evaluated as +1 nucleophiles (cyan) and the native cyclization sites (grey). (B) Schematic representation of the corresponding SIML expression constructs generated using the split Gp41-1 intein, illustrating insertion of the bacteriocin sequence between the N- and C-terminal intein fragments at each selected serine position. (C) Antimicrobial activity of IV-CFPS/SIML products assessed by spot-on-agar assays against *P. damnosus* CECT 4797. Samples were tested using serial two-fold dilutions ranging from 1:2 to 1:65536.

GarML produced using the IV-CFPS/SIML platform displayed relatively modest yet consistent antimicrobial activity across all three ligation sites tested. Among these, Gp41-1-GarML_S29 (dilution 1: 32) and Gp41-1-GarML_S32 (dilution 1:64) exhibited the strongest inhibitory activity, whereas Gp41-1-GarML_S19 (dilution 1:16) showed the lowest activity. The predicted α-helical three-dimensional structure of GarML, together with the distribution of neighboring residues, may facilitate efficient intein-mediated splicing and recovery of an active conformation across multiple junction positions. In contrast, AmCyc exhibited a pronounced dependence on the cyclization site. Only the mature bacteriocin cyclized at S23 (Gp41-1-ACC_S23; dilution 1:8192) displayed high antimicrobial activity, whereas constructs cyclized at S3 (Gp41-1-ACC_S3; dilution 1:8), S8 (Gp41-1-ACC_S8; dilution 1:64), or S30 (Gp41-1-ACC_S30; dilution 1:8) showed substantially weaker inhibition. These results suggest that AmCyc requires a more stringent cyclization geometry, likely reflecting peptide-specific structural constraints associated with helix boundaries or loop connectivity that limit the number of tolerated ligation sites.

For GarML the relative low antimicrobial activity observed across all constructs did not correlate with the canonical flanking-sequence preferences described for Gp41-1, which typically favors flexible residues at the −1/+1 positions [17,20]. Notably, the most active variant, GarML_S32 (−1/+1 = F/G), would be predicted to be less favorable than GarML_S19 (−1/+1 = V/A) based on these criteria. This discrepancy underscores the importance of higher-order structural context, such as helix packing and long-range intramolecular interactions, in overriding primary sequence preferences when determining intein splicing efficiency and functional peptide activity. Similar context-dependent effects have been reported for engineered and evolved split inteins, in which global structural compatibility, rather than local extein sequence, governs reaction kinetics and yield [41,42].

Collectively, these findings indicate that the positional effects observed for EntAS-48 are not universally conserved among circular bacteriocins. Whereas some peptides, such as GarML, tolerate shifts in the ligation site, others, including AmCyc, exhibit a strict dependence on precise junction placement. These results further emphasize that split intein-mediated cyclization outcomes cannot be reliably predicted from sequence motifs alone and highlight the necessity of empirically evaluating multiple ligation sites when applying the IV-CFPS/SIML platform to newly characterized or putative circular bacteriocins.

### 3.4. Functional Analysis of Circular Bacteriocin Variants Produced via the Gp41-1 Split Intein and the IV-CFPS/SIML Platform

After identifying the optimal intein–cyclization site combinations for IV-CFPS/SIML-derived production of EntAS-48, GarML, and AmCyc, we next examined whether these standardized conditions could be applied to systematically evaluate the production and antimicrobial activity of naturally occurring circular bacteriocin homologs. Using the split Gp41-1 intein and the optimal conserved cyclization sites for each scaffold (S41 for EntAS-48, S29 for GarML, and S23 for AmCyc), we generated and screened a panel of 27 circular bacteriocin variants (Fig. 5, Table S3). For each bacteriocin family, the selected serine residue was conserved across all homologs, enabling the use of a common cyclization site without introducing nonnative mutations (Fig. 5). Although Gp41-1-GarML_S32 showed slightly higher activity in preliminary assays (Fig. 4), serine S29 was selected for GarML-derived variant construction because it was conserved in all homologs, whereas only two of the five variants encoded a serine at position 32.

**Fig. 5.**
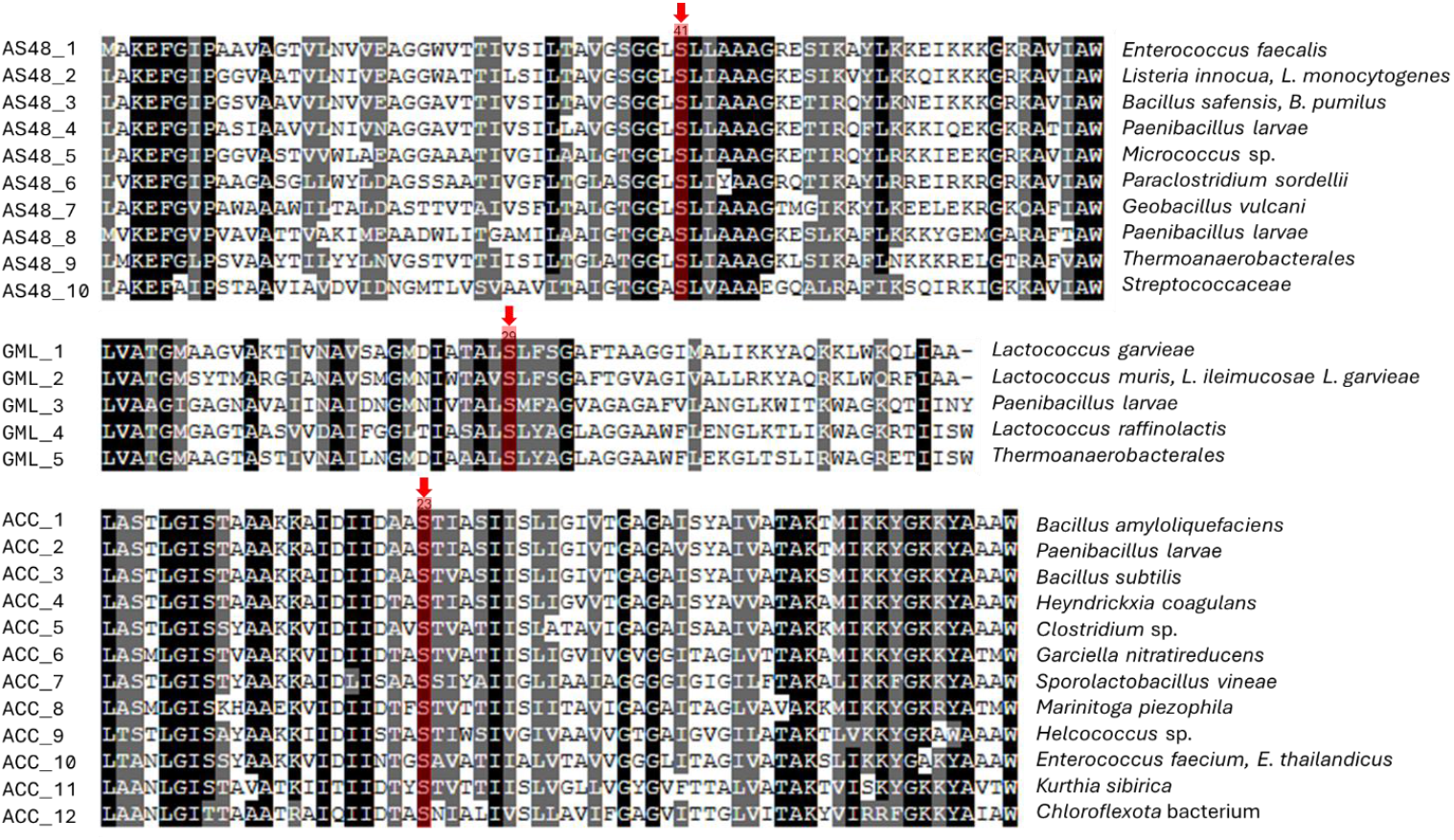
Multiple sequence alignments of EntAS-48 (AS48), GarML (GML) and AmCyc (ACC) variants generated using Color Align Conservation. Identical residues are indicated by a black background, similar residues by a gray background, and nonconserved residues by a white background. Red arrows denote the conserved serine residue selected as the +1 nucleophile for intein-mediated cyclization within each bacteriocin group.

Twenty-seven PCR-assembled SIML expression cassettes were used as DNA templates for IV-CFPS, and the resulting bacteriocin variants were evaluated against a panel of Gram-positive and Gram-negative indicator strains (Table 1). This panel included four previously characterized circular bacteriocins EntAS-48 (AS48_1), GarML (GML_1), AmCyc (ACC_1), and enterocin NKR-5-3B (ACC_10), together with nine native EntAS-48 variants (AS48_2 to AS48_10), four GarML variants (GML_2 to GML_5), and ten AmCyc variants (ACC_2 to ACC_9, ACC_11, and ACC_12). The analyzed sequences originated from microorganisms spanning broad phylogenetic and ecological diversity, including lactic acid bacteria (LAB) (e.g. *Lactococcus raffinolactis, L. ileimucosae, L. garvieae*), environmental and spore-forming Firmicutes (*Bacillus safensis, B. pumilus, Sporolactobacillus vineae*), insect pathogens (*Paenibacillus larvae*), anaerobic and clostridia-related taxa (*Paraclostridium sordellii, Garciella nitratireducens, Clostridium* spp.), and thermophilic or deep-branching lineages such as *Thermoanaerobacterales, Geobacillus vulcani, Marinitoga piezophila*, several *Chloroflexota* representatives, and others (Table S3). This diversity indicates that circular bacteriocins are not confined to classical LAB or *Bacillus* spp., but are widely distributed across environmental, host-associated, anaerobic, and extremophilic ecosystems. The occurrence of circular bacteriocins in extremophilic taxa further suggests that head-to-tail cyclization may confer an adaptive advantage under harsh physicochemical conditions, where enhanced peptide stability is critical for antimicrobial activity [15].

**Table 1.**
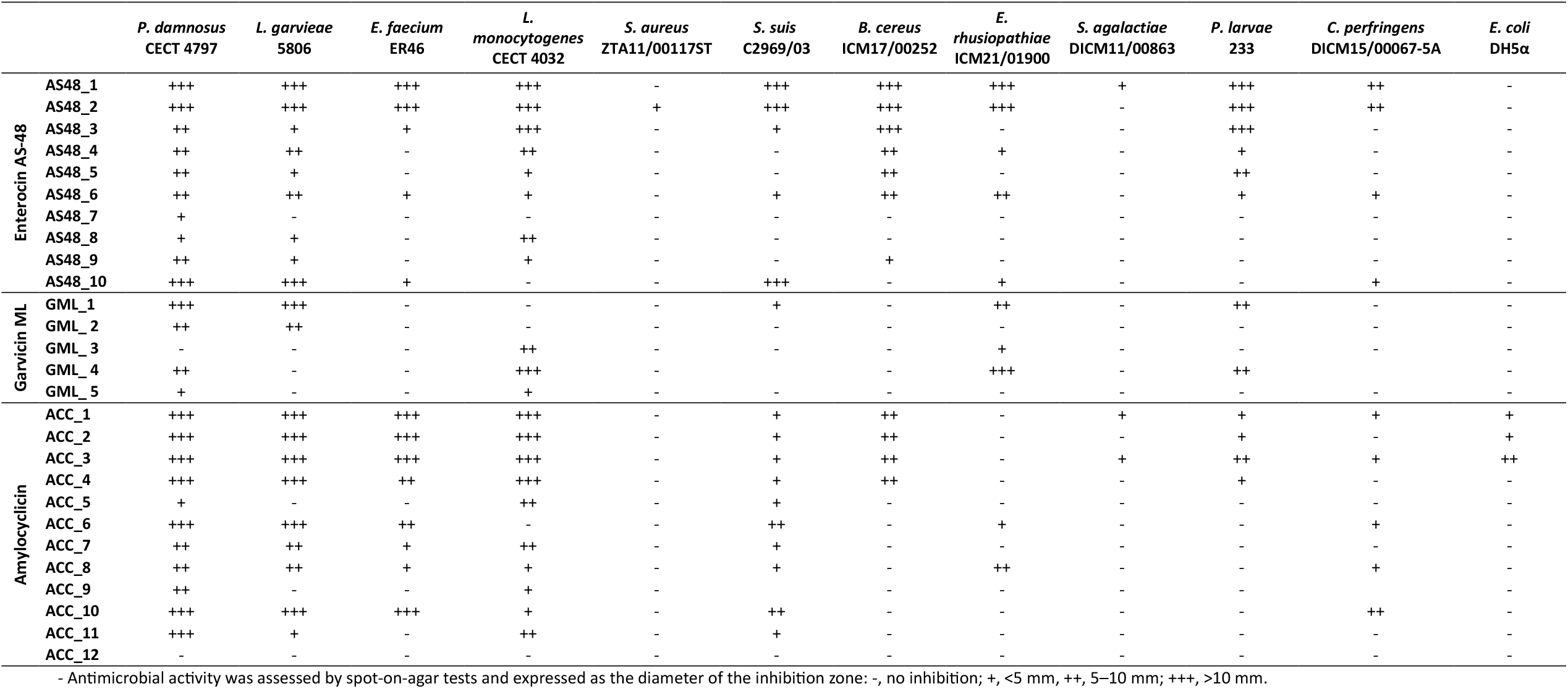
Antimicrobial activity of circular bacteriocin variants against selected indicator bacteria.

Across all bacteriocin variant groups, substantial variation in antimicrobial activity and target range was observed, indicating that even minor sequence differences can markedly influence antimicrobial potency and specificity. The four reference bacteriocins exhibited a strong and broad inhibitory activity against multiple Gram-positive species, thereby validating the reliability of the IV-CFPS/SIML platform for the production and functional evaluation of circular bacteriocins. Most AS48 variants displayed activity primarily against Gram-positive bacteria. Among them, the AS48_1 and AS48_2 variants showed the broadest inhibition spectra, with activity against *P. damnosus, Lactococcus garvieae, Enterococcus faecium, Listeria monocytogenes, Streptococcus suis, Bacillus cereus, Erysipelothrix rhusiopathiae, Paenibacillus larvae*, and *Clostridium perfringens*. In contrast, other variants, such as AS48_3 and AS48_10, exhibited narrower activity profiles. None of the AS48 variants exhibited detectable inhibitory activity against *E. coli*.

The GarML variants inhibited exclusively Gram-positive bacteria. Among these, GML_1 exhibited the highest antimicrobial potency, whereas GML_5 showed the weakest activity, with intermediate inhibition observed for GML_2–GML_4. In contrast, the AmCyc homologs displayed the greatest functional diversity. Several variants (ACC_1– ACC_4) showed strong antimicrobial activity against a broad range of Gram-positive strains, and notably, ACC_1 and ACC_3 also inhibited *E. coli*. Among all variants tested, ACC_3 demonstrated the highest overall antimicrobial potency, whereas ACC_12 showed no detectable activity. These findings suggest that specific structural or sequence features of the AmCyc scaffold may facilitate interactions with Gram-negative outer membranes or alternative cellular targets, thereby expanding its antimicrobial spectrum.

Notably, many bacteriocin variants were identified in genomes from ecologically diverse and often inaccessible bacterial lineages, including extremophiles and spore-forming taxa adapted to severe environmental stress. Among these, ACC_11 was particularly noteworthy, as it derives from an *AmCyc*-like gene identified in *Kurthia sibirica* isolated from the large intestine of a mammoth preserved in East Siberian permafrost. Although ACC_11 exhibited only moderate antimicrobial activity, its successful *in vitro* expression and functional validation demonstrate that the SIML platform can recover and evaluate putative circular bacteriocins from rare, unconventional, or even extinct ecological niches. Collectively, these findings highlight the capacity of the IV-CFPS/SIML platform to translate genomic predictions into experimentally validated antimicrobial peptides and to substantially expand the repertoire of circular bacteriocins, including those from extreme or unconventional environments.

### 3.5. *In Vivo* Validation of Optimal Cyclization Site-Split Intein Combinations for the Production of EntAS-48 (AS48_1) and AmCyc variant 3 (ACC_3)

To determine whether SIML expression cassettes also support *in vivo* production and functional expression of circular bacteriocins, we evaluated whether the optimal cyclization site-split intein combinations identified using the IV-CFPS/SIML platform could be translated to recombinant *E. coli*. Specifically, we examined the *in vivo* production of EntAS-48 (AS48_1) and AmCyc variant 3 (ACC_3). GarML was not included, as its SIML-mediated production in *E. coli* has been previously demonstrated [14]. Both AS48_1 and ACC_3 were produced in *E. coli* BL21 (DE3) harboring pCir-Gp41-1-AS48_1 (S41) and pCir-Gp41-ACC_3 (S23), respectively. Bacteriocins were purified from cellular-soluble fractions by hydrophobic interaction chromatography (HIC) followed by reverse-phase fast protein liquid chromatography (RP-FPLC). Fractions exhibiting the highest antimicrobial activity were analyzed by MALDI–TOF MS. Purified AS48_1 showed an observed molecular mass of 7,149.45 Da, identical to that of the native circular peptide produced by *E. faecalis* [34,43], and 18 Da lower than its calculated linear precursor (7,163.08 Da). Similarly, ACC_3 exhibited an observed molecular mass of 6,352.640 Da, likewise about 18 Da lower than its calculated linear precursor (6,371.57 Da), consistent with water loss during head-to-tail cyclization (Fig. 6). Collectively, these results demonstrate that cyclization site-split intein combinations optimized *in vitro* using the IV-CFPS/SIML platform can be directly translated to efficient *in vivo* production of circular bacteriocins in recombinant *E. coli*.

**Fig. 6.**
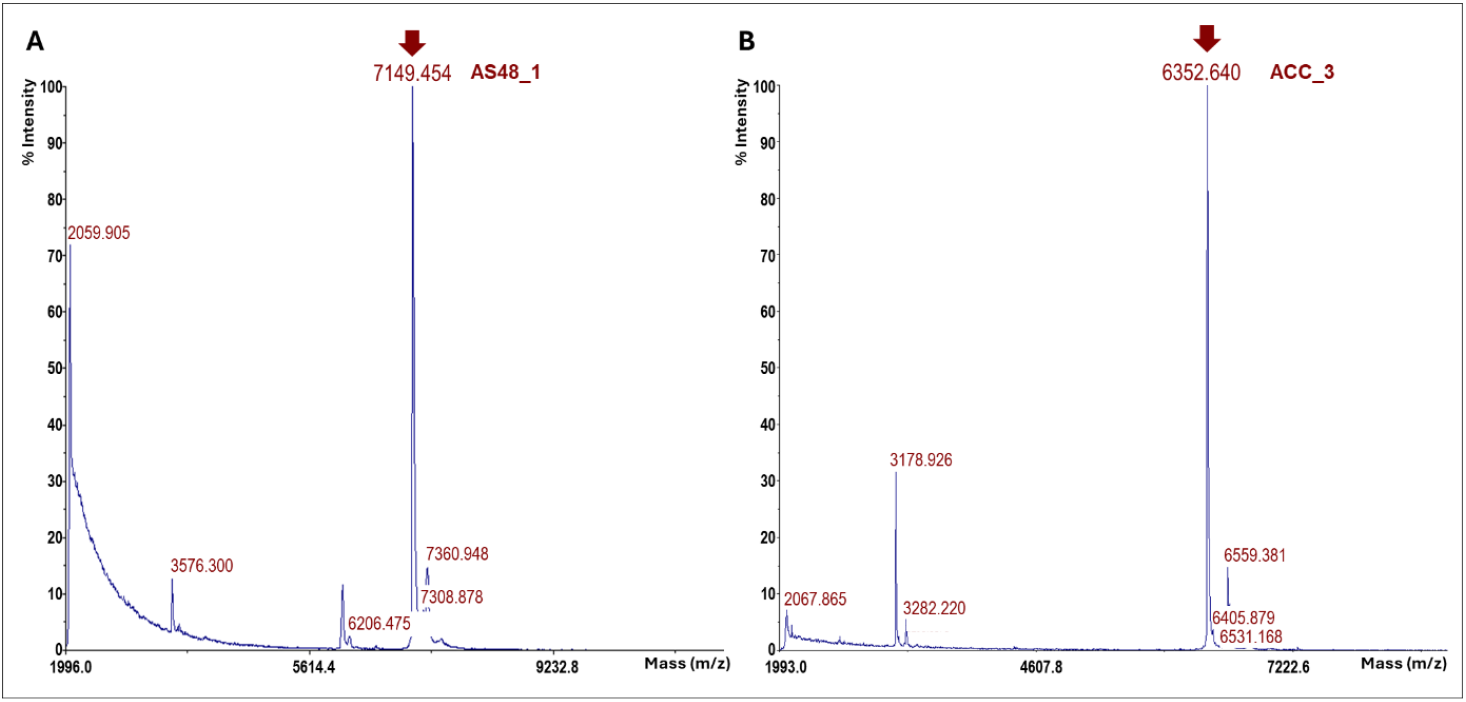
MALDI-TOF MS of the RP-FPLC–purified active fractions obtained from *E. coli* BL21 (DE3) transformed with pCir-Gp41-1-AS48_1 (S41) (A), and pCir-Gp41-1-ACC_3 (S23). Spectra were acquired in linear mode using sinapinic acid as the matrix. Maroon arrows indicate peptide peaks corresponding to the observed molecular masses of EntAS-48 (AS-48_1) and AmCyc variant 3 (ACC_3).

## 4. Conclusions

The accelerating emergence of antibiotic-resistant pathogens underscores the urgent need for synthetic biology platforms capable of rapidly discovering, validating, and engineering new antimicrobial agents. Circular bacteriocins are particularly attractive due to their exceptional structural stability and potent antimicrobial activity. However, their broader exploration has been constrained by the complexity of native biosynthetic pathways, limiting mechanistic insight and heterologous production. Here, we present a versatile synthetic-biology framework that integrates *in vitro* cell-free protein synthesis with Split Intein-Mediated Ligation (IV-CFPS/SIML), enabling efficient production, head-to-tail cyclization, and functional expression of mature circular bacteriocins without the need for additional gene cluster-encoded proteins involved in processing, transport, regulation or immunity.

Systematic evaluation of cyclization-site selection and split intein identity revealed that both parameters strongly influence IV-CFPS/SIML production efficiency and antimicrobial activity of enterocin AS-48, garvicin ML, and amylocyclicin, with optimal cyclization sites combined with the split Gp41-1 intein consistently yielding superior activity. These conditions enabled functional production of 27 naturally occurring bacteriocin variants, uncovering substantial diversity in antimicrobial potency and target range and demonstrating that even minor sequence variations can markedly alter biological activity. Collectively, these findings establish the IV-CFPS/SIML platform as a rapid and scalable approach for the production, screening, and functional expression of circular bacteriocins, including those derived from unconventional or difficult-to-culture microorganisms, thereby expanding access to circular antimicrobial peptide diversity.

Importantly, transfer of SIML constructs to pUC-derived vectors enabled *in vivo* production, cyclization, and functional expression of selected bacteriocins in recombinant *E. coli*, demonstrating direct transferability of optimized IV-CFPS/SIML designs to cellular systems. Collectively, these findings establish SIML as a rapid, modular, and scalable platform for *in vitro* and *in vivo* production, screening, and engineering of circular bacteriocins and highlight split intein-mediated cyclization as a powerful strategy for expanding antimicrobial peptide diversity and guiding next-generation peptide design.

## Associated content

### Data Availability Statement

The data underlying this article are available in the article and in its online Supporting Information.

### Supporting information

The Supporting Information is available free of charge at XXXX

**Fig. S1**. Schematic representation of SIML expression cassette assembly by overlap-extension PCR. The SIML constructs were generated using a three-fragment assembly strategy. Fragment A contains the T7 promoter followed by the C-terminal fragment of the split intein (I_C_). Fragment B encodes the mature circular bacteriocin sequence, beginning at the selected serine residue used as the +1 nucleophile for cyclization, and is flanked by regions homologous to I_C_ at the 5′ end and to the N-terminal intein fragment (I_N_) at the 3′ end. Fragment C contains the I_N_ fragment followed by a transcription terminator. The three fragments were assembled by overlap-extension PCR to generate a complete T7–I_C_–bacteriocin–I_N_–terminator expression cassette, which was purified and used directly as a DNA template for *in vitro* cell-free protein synthesis (IV-CFPS) reactions within the SIML framework. **Table S1**. Nucleotide and amino acid sequences used for SIML construct design and circular bacteriocin variant libraries. The table summarizes the DNA and protein sequences employed in the generation of SIML expression cassettes. The upper section lists the nucleotide sequences of the T7 promoter, transcription terminator, and primers used for cassette assembly. The middle section shows the amino acid sequences of the split inteins used in this study, including NpuDnaE, Gp41-1, and SspGyrB, with their corresponding C-terminal (I_C_) and N-terminal (I_N_) fragments. The lower section details the amino acid sequences of the circular bacteriocins and their variants evaluated. **Table S2**. Origin, growth media, and incubation conditions of the bacterial strains used in this study. **Table S3**. Amino acid sequences and physicochemical properties of the circular bacteriocin variants analyzed in this study.

## Author Information

### Authors

**Ester Sevillano –** *Departamento de Nutrición y Ciencia de los Alimentos (NUTRYCIAL), Sección Departamental de Nutrición y Ciencia de los Alimentos (SD-NUTRYCIAL), Facultad de Veterinaria, Universidad Complutense de Madrid (UCM), Avenida Puerta de Hierro, s/n, 28040 Madrid, Spain*.

**Mohamed El Bakkoury** - *Syngulon SA, Seraing, Belgium*.

**Irene Lafuente -** *Departamento de Nutrición y Ciencia de los Alimentos (NUTRYCIAL), Sección Departamental de Nutrición y Ciencia de los Alimentos (SD-NUTRYCIAL), Facultad de Veterinaria, Universidad Complutense de Madrid (UCM), Avenida Puerta de Hierro, s/n, 28040 Madrid, Spain*.

**Nuria Peña -** *Departamento de Nutrición y Ciencia de los Alimentos (NUTRYCIAL), Sección Departamental de Nutrición y Ciencia de los Alimentos (SD-NUTRYCIAL), Facultad de Veterinaria, Universidad Complutense de Madrid (UCM), Avenida Puerta de Hierro, s/n, 28040 Madrid, Spain*.

**Luis M. Cintas -** *Departamento de Nutrición y Ciencia de los Alimentos (NUTRYCIAL), Sección Departamental de Nutrición y Ciencia de los Alimentos (SD-NUTRYCIAL), Facultad de Veterinaria, Universidad Complutense de Madrid (UCM), Avenida Puerta de Hierro, s/n, 28040 Madrid, Spain*.

**Philippe Gabant** - *Syngulon SA, Seraing, Belgium*.

**Pablo E. Hernández -** *Departamento de Nutrición y Ciencia de los Alimentos (NUTRYCIAL), Sección Departamental de Nutrición y Ciencia de los Alimentos (SD-NUTRYCIAL), Facultad de Veterinaria, Universidad Complutense de Madrid (UCM), Avenida Puerta de Hierro, s/n, 28040 Madrid, Spain*.

### Author Contributions

E.M.-A., P.G., P.E.H. and J.B. conceptualized the study. E.S., M.E.B., P.G. and J.B. designed the plasmids and gene constructs. E.S., I.L., N.P. and M.E.B. performed the experiments. E.S., M.E.B., P.G., P.E.H., and J.B. analyzed the data.E.S. and J.B. generated the figures. E.S. wrote the manuscript with input from all authors. P.G. P.E.H. and J.B. reviewed and edited the manuscript. E.M.-A., L.M.C., P.G., P.E.H. and J.B. supervised the research. E.M.-A., L.M.C., P.G., P.E.H. and J.B. secured funding for the study.

### Notes

The authors declare the following competing financial interests: P.G. and J.B. are inventors on a pending patent application entitled “*Bacteriocin polypeptides, nucleic acids encoding same, and methods of use thereof*.” All remaining authors declare no competing financial interest.

## Acknowledgements

The authors thank the Proteomics Unit of the Universidad Complutense de Madrid (UCM), for technical assistance with proteomic analyses.

