## Supplementary material for "An Enhanced Split Intein-Mediated Ligation (SIML) Platform for Rapid Discovery and Functional Screening of Circular Bacteriocins": Supplemetary Information

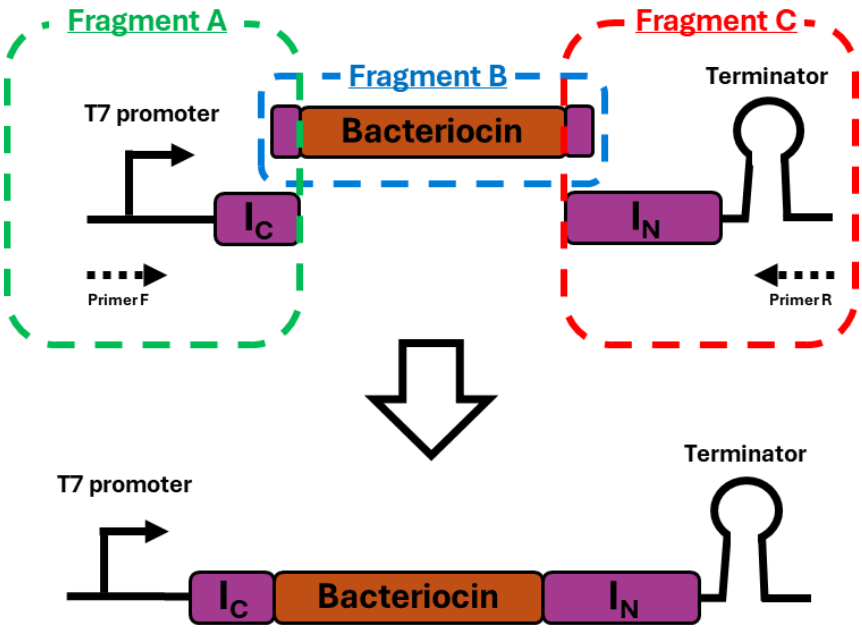


**Fig. S1. Schematic representation of SIML expression cassette assembly by overlap-extension PCR.** The SIML constructs were generated using a three-fragment assembly strategy. **Fragment A** contains the T7 promoter followed by the C-terminal fragment of the split intein ($I_C$). **Fragment B** encodes the mature circular bacteriocin sequence, beginning at the selected serine residue used as the +1 nucleophile for cyclization, and is flanked by regions homologous to $I_C$ at the 5′ end and to the N-terminal intein fragment ($I_N$) at the 3′ end. **Fragment C** contains the $I_N$ fragment followed by a transcription terminator. The three fragments were assembled by overlap-extension PCR to generate a complete T7–$I_C$–bacteriocin–$I_N$–terminator expression cassette, which was purified and used directly as a DNA template for *in vitro* cell-free protein synthesis (IV-CFPS) reactions within the SIML framework.

**Table S1. Nucleotide and amino acid sequences used for SIML construct design and circular bacteriocin variant libraries.** The table summarizes the DNA and protein sequences employed in the generation of SIML expression cassettes. The upper section lists the nucleotide sequences of the T7 promoter, transcription terminator, and primers used for cassette assembly. The middle section shows the amino acid sequences of the split inteins used in this study, including NpuDnaE, Gp41-1, and SspGyrB, with their corresponding C-terminal ($I_C$) and N-terminal ($I_N$) fragments. The lower section details the amino acid sequences of the circular bacteriocins and their variants evaluated.

| | **Nucleotidic sequence (5´- 3´)** |
|---|---|
| **T7 promoter** | GCGAATTAATACGACTCACTATAGGGCTTAAGTATAAGGAGGAAAAAAT |
| **Terminator** | TAACTAGCATAACCCCTCTCTAAACGGAGGGGTTT |
| **Primer F** | GCGAATTAATACGACTCACTATAG |
| **Primer R** | AAACCCCTCCGTTTAGAG |

| **Split intein** | **Amino acid sequence** |
|---|---|
| **NpuDnaE $I_C$** | MIKIATRKYLGKQNVYDIGVERYHNFALKNGFIASN |
| **NpuDnaE $I_N$** | CLSYDTEILTVEYGILPIGKIVEKRIECTVYSVDNNGNIYTQPVAQWHDRGEQEVFEYCLEDGCLIRATK DHKFMTVDGQMMPIDEIFERELDLMRVDNLPNGT |
| **Gp41-1 $I_C$** | MMLKKILKIEELDERELIDIEVSGNHLFYANDILTHN |
| **Gp41-1 $I_N$** | CLDLKTQVQTPQGMKEISNIQVGDLVLSNTGYNEVLNVFPKSKKKSYKITLEDGKEIICSEEHLFPTQTG EMNISGGLKEGMCLYVKE |
| **SspGyrB $I_C$** | MEAVLNYNHRIVNIEAVSETIDVYDIEVPHTHNFALASGVFVHN |
| **SspGyrB $I_N$** | CFSGDTLVALTDGRSVSFEQLVEEEKQGKQNFCYTIRHDGSIGVEKIINARKTKTNAKVIKVTLDNGESII CTPDHKFMLRDGSYKCAMDLTLDDSLMPLHRKISTTEDSGHA |

| **Bacteriocin** | **Amino acid sequence** |
|---|---|
| **AS48_S30** | SILTAVGSGGLSLLAAAGRESIKAYLKKEIKKKGKRAVIAWMAKEFGIPAAVAGTVLNVVEAGGWVTTIV |
| **AS48_S37** | SGGLSLLAAAGRESIKAYLKKEIKKKGKRAVIAWMAKEFGIPAAVAGTVLNVVEAGGWVTTIVSILTAVG |
| **AS48_S41 (AS48_1)** | SLLAAAGRESIKAYLKKEIKKKGKRAVIAWMAKEFGIPAAVAGTVLNVVEAGGWVTTIVSILTAVGSGGL |
| **AS48_S50** | SIKAYLKKEIKKKGKRAVIAWMAKEFGIPAAVAGTVLNVVEAGGWVTTIVSILTAVGSGGLSLLAAAGRE |
| **GarML_S19** | SAGMDIATALSLFSGAFTAAGGIMALIKKYAQKKLWKQLIAALVATGMAAGVAKTIVNAV |
| **GarML_S29 (GarML_1)** | SLFSGAFTAAGGIMALIKKYAQKKLWKQLIAALVATGMAAGVAKTIVNAVSAGMDIATAL |
| **GarML_S32** | SGAFTAAGGIMALIKKYAQKKLWKQLIAALVATGMAAGVAKTIVNAVSAGMDIATALSLF |
| **ACC_S3** | STLGISTAAAKKAIDIIDAASTIASIISLIGIVTGAGAISYAIVATAKTMIKKYGKKYAAAWLA |
| **ACC_S8** | STAAAKKAIDIIDAASTIASIISLIGIVTGAGAISYAIVATAKTMIKKYGKKYAAAWLASTLGI |
| **ACC_S23 (ACC_1)** | STIASIISLIGIVTGAGAISYAIVATAKTMIKKYGKKYAAAWLASTLGISTAAAKKAIDIIDAA |
| **ACC_S30** | SLIGIVTGAGAISYAIVATAKTMIKKYGKKYAAAWLASTLGISTAAAKKAIDIIDAASTIASII |
| **AS48_2** | SLIAAAGKESIKVYLKKQIKKKGRKAVIAWLAKEFGIPGGVAATVLNIVEAGGWATTILSILTAVGSGGL |
| **AS48_3** | SLIAAAGKETIRQYLKNEIKKKGRKAVIAWLAKEFGIPGSVAAVVLNVVEAGGAVTTIVSILTAVGSGGL |
| **AS48_4** | SLLAAAGKETIRQFLKKKIQEKGKRATIAWLAKEFGIPASIAAVVLNIVNAGGAVTTIVSILLAVGSGGL |
| **AS48_5** | SLIAAAGKETIRQYLRKKIEEKGRKAVIAWLAKEFGIPGGVASTVVWLAEAGGAAATIVGILAALGTGGL |
| **AS48_6** | SLIYAAGRQTIKAYLRREIRKRGRKAVIAWLVKEFGIPAAGASGLLWYLDAGSSAATIVGFLTGLASGGL |
| **AS48_7** | SLIAAAGTMGIKKYLKEELEKRGKQAFIAWLAKEFGVPAWAAAWILTALDASTTVTAIVSFLTALGTGGL |
| **AS48_8** | SLLAAAGKESLKAFLKKKYGEMGARAFTAWMVKEFGVPVAVATTVAKIMEAADWLITGAMILAAIGTGGA |
| **AS48_9** | SLIAAAGKLSIKAFLNKKKRELGTRAFVAWLMKEFGLPSVAAYTILYYLNVGSTVTTIISILTGLATGGL |
| **AS48_10** | SLVAAAEGQALRAFIKSQIRKIGKKAVIAWLAKEFAIPSTAAVIAVDVIDNGMTLVSVAAVITAIGTGGA |
| **GarML_2** | SLFSGAFTGVAGIVALLRKYAQRKLWQRFIAALVATGMSYTMARGIANAVSMGMNIWTAV |

| | |
|---|---|
| **GarML_3** | SMFAGVAGAGAFVLANGLKWITKWAGKQTIINYLVAAGIGAGNAVAIINAIDNGMNIVTAL |
| **GarML_4** | SLYAGLAGGAAWFLENGLKTLIKWAGKRTIISWLVATGMGAGTAASVVDAIFGGLTIASAL |
| **GarML_5** | SLYAGLAGGAAWFLEKGLTSLIRWAGRETIISWLVATGMAAGTASTIVNAILNGMDIAAAL |
| **ACC_2** | STIASIISLIGIVTGAGAVSYAIVATAKTMIKKYGKKYAAAWLASTLGISTAAAKKAIDIIDAA |
| **ACC_3** | STVASIISLIGIVTGAGAISYAIVATAKSMIKKYGKKYAAAWLASTLGISTAAAKKAIDIIDAA |
| **ACC_4** | STIASIISLIGVVTGAGAISYAVVATAKAMIKKYGKKYAAAWLASTLGISTAAAKKAIDIIDTA |
| **ACC_5** | STVATIISLATAVIGAGAISAAIVATAKKMIKKYGKKYAAAWLASTLGISSYAAKKVIDIIDAV |
| **ACC_6** | STVATIISLIGVIVGVGGITAGLVTTAKAMIKKYGKKYATMWLASMLGISTVAAKKVIDIIDTA |
| **ACC_7** | SSIYAIIGLIAAIAGGGGIGIGILFTAKALIKKFGKKYAAAWLASTLGISTYAAKKAIDLISAA |
| **ACC_8** | STVTTIISIITAVIGAGAITAGLVAVAKKMIKKYGKRYATMWLASMLGISKHAAEKVIDIIDTF |
| **ACC_9** | STIWSIVGIVAAVVGTGAIGVGILATAKTLVKKYGKAWAAAWLTSTLGISAYAAKKIIDIISTA |
| **ACC_10** | SAVATIIALVTAVVGGGLITAGIVATAKSLIKKYGAKYAAAWLTANLGISSYAAKKVIDIINTG |
| **ACC_11** | STVTTIISLVGLLVGYGVFTTALVATAKTVISKYGKKYAVTWLAANLGISTAVATKIITIIDTY |
| **ACC_12** | SNIALIVSLLAVIFGAGVITTGLVITAKYVIRRFGKKYAIAWLAANLGITTAAATRAIQIIDTA |

**Table S2.** Origin, growth media, and incubation conditions of the bacterial strains used in this study.

| Strain[a] | Origin[b] | Growth medium | Incubation conditions |
|---|---|---|---|
| *Pediococcus damnosus* CECT 4797 | CECT | MRS | 32 °C/ Aerobiosis |
| *Lactococcus garvieae* 5806 | DNBTA | MRS | 32 °C/ Aerobiosis |
| *Enterococcus faecium* ER46 | VISAVET | MRS | 37 °C/ Aerobiosis |
| *Listeria monocytogenes* CECT 4032 | CECT | BHI | 37 °C/ Aerobiosis |
| *Staphylococcus aureus* ZTA11/00117ST | VISAVET | BHI | 37 °C/ Aerobiosis |
| *Streptococcus suis* C2969/03 | VISAVET | BHI | 37 °C/ Aerobiosis |
| *Bacillus cereus* ICM17/00252 | VISAVET | BHI | 37 °C/ Aerobiosis |
| *Erysipelothrix rhusiopathiae* ICM21/01900 | VISAVET | BHI | 37 °C/ Aerobiosis |
| *Streptococcus agalactiae* DICM11/00863 | VISAVET | BHI | 37 °C/ Aerobiosis |
| *Paenibacillus larvae* DB25 | DNBTA | BHI | 37 °C/ Aerobiosis |
| *Clostridium perfringens* DICM15/00067-5A | VISAVET | BHI | 37 °C/ Anaerobiosis |
| *Escherichia coli* DH5α | Thermofisher | LB | 37 °C/Agitation |
| *Escherichia coli* BL21 (DE3) | Thermofisher | LB | 37 °C/Agitation |

[a] Strains were used as indicator microorganisms, with the exception of *E. coli* BL21 (DE3), which was used for the *in vivo* production and purification of circular bacteriocins.

[b] DNBTA: Departamento de Nutrición, Bromatología y Tecnología de los Alimentos, Facultad de Veterinaria, Universidad Complutense de Madrid (UCM), Madrid, (Spain). VISAVET: Centro de Vigilancia Sanitaria Veterinaria, Universidad Complutense de Madrid (UCM), Madrid, (Spain). CECT: Colección Española de Cultivos Tipo, Valencia, (Spain).

**Table S3**. Amino acid sequences and physicochemical properties of the circular bacteriocin variants analyzed in this study.

| Bacteriocin variant | Species origin | Percent Identity[a] | Length (aa) | Molecular mass (Da) | Theoretical pI | Net charge | Aliphatic index | GRAVY[b] | Mature amino acid sequence |
|---|---|---|---|---|---|---|---|---|---|
| AS-48_1 | *Enterococcus faecalis* | 100% | 70 | 7150 | 10.09 | 6 | 117.14 | 0.539 | MAKEFGIPAAVAGTVLNVVEAGGWVTTIVSILTAVGSGGLSLLAAAGRESIKAYLKKEIKKKGKRAVIAW |
| AS-48_2 | *Listeria innocua, L. monocytogenes* | 81% | 70 | 7117 | 10.17 | 7 | 124.14 | 0.551 | LAKEFGIPGGVAATVLNIVEAGGWATTILSILTAVGSGGLSLIAAAGKESIKVYLKKQIKKKGRKAVIAW |
| AS-48_3 | *Bacillus safensis, B. pumilus* | 80% | 70 | 7087 | 10.00 | 5 | 125.43 | 0.579 | LAKEFGIPGSVAAVVLNVVEAGGAVTTIVSILTAVGSGGLSLIAAAGKETIRQYLKNEIKKKGRKAVIAW |
| AS-48_4 | *Paenibacillus larvae* | 74% | 70 | 7127 | 10.38 | 6 | 131.14 | 0.661 | LAKEFGIPASIAAVVLNIVNAGGAVTTIVSILLAVGSGGLSLLAAAGKETIRQFLKKKIQEKGKRATIAW |
| AS-48_5 | *Micrococcus* sp. | 63% | 70 | 7072 | 9.87 | 4 | 118.71 | 0.510 | LAKEFGIPGGVASTVVWLAEAGGAAATIVGILAALGTGGLSLIAAAGKETIRQYLRKKIEEKGRKAVIAW |
| AS-48_6 | *Paraclostridium sordellii* | 59% | 70 | 7362 | 10.66 | 7 | 113.14 | 0.386 | LVKEFGIPAAGASGLLWYLDAGSSAATIVGFLTGLASGGLSLIYAAGRQTIKAYLRREIRKRGRKAVIAW |
| AS-48_7 | *Geobacillus vulcani* | 59% | 70 | 7322 | 9.23 | 2 | 110.43 | 0.574 | LAKEFGVPAWAAAWILTALDASTTVTAIVSFLTALGTGGLSLIAAAGTMGIKKYLKEELEKRGKQAFIAW |
| AS-48_8 | *Paenibacillus larvae* | 57% | 70 | 7185 | 9.52 | 3 | 99.29 | 0.633 | MVKEFGVPVAVATTVAKIMEAADWLITGAMILAAIGTGGASLLAAAGKESLKAFLKKKYGEMGARAFTAW |
| AS-48_9 | *Thermoanaerobacterales* | 53% | 70 | 7416 | 10.09 | 6 | 124.14 | 0.680 | LMKEFGLPSVAAYTILYYLNVGSTVTTIISILTGLATGGLSLIAAAGKLSIKAFLNKKKRELGTRAFVAW |
| AS-48_10 | *Streptococcaceae* | 43% | 70 | 7017 | 9.82 | 3 | 128.43 | 0.881 | LAKEFAIPSTAAVIAVDVIDNGMTLVSVAAVITAIGTGGASLVAAAEGQALRAFIKSQIRKIGKKAVIAW |
| GML_1 | *Lactococcus garvieae* | 100% | 60 | 6007 | 10.13 | 5 | 115.83 | 0.887 | LVATGMAAGVAKTIVNAVSAGMDIATALSLFSGAFTAAGGIMALIKKYAQKKLWKQLIAA |
| GML_ 2 | *L. muris L. ileimucosae L. garvieae* | 65% | 60 | 6422 | 11.58 | 6 | 101.00 | 0.687 | LVATGMSYTMARGIANAVSMGMNIWTAVSLFSGAFTGVAGIVALLRKYAQRKLWQRFIAA |
| GML_ 3 | *Paenibacillus larvae* | 47% | 61 | 6086 | 9.53 | 2 | 121.80 | 0.913 | LVAAGIGAGNAVAIINAIDNGMNIVTALSMFAGVAGAGAFVLANGLKWITKWAGKQTIINY |
| GML_ 4 | *Lactococcus raffinolactis* | 46% | 61 | 6092 | 9.53 | 2 | 117.05 | 0.872 | LVATGMGAGTAASVVDAIFGGLTIASALSLYAGLAGGAAWFLENGLKTLIKWAGKRTIISW |
| GML_ 5 | *Thermoanaerobacterales* | 76% | 61 | 6190 | 6.18 | 0 | 120.33 | 0.844 | LVATGMAAGTASTIVNAILNGMDIAAALSLYAGLAGGAAWFLEKGLTSLIRWAGRETIISW |
| ACC_1 | *Bacillus amyloliquefaciens* | 100% | 64 | 6382 | 9.82 | 5 | 125.47 | 0.850 | LASTLGISTAAAKKAIDIIDAASTIASIISLIGIVTGAGAISYAIVATAKTMIKKYGKKYAAAW |
| ACC_2 | *Paenibacillus larvae* | 98% | 64 | 6368 | 9.82 | 5 | 123.91 | 0.845 | LASTLGISTAAAKKAIDIIDAASTIASIISLIGIVTGAGAVSYAIVATAKTMIKKYGKKYAAAW |
| ACC_3 | *Bacillus subtilis* | 97% | 64 | 6354 | 9.82 | 5 | 123.91 | 0.844 | LASTLGISTAAAKKAIDIIDAASTVASIISLIGIVTGAGAISYAIVATAKSMIKKYGKKYAAAW |
| ACC_4 | *Heyndrickxia coagulans* | 94% | 64 | 6354 | 9.82 | 5 | 122.34 | 0.841 | LASTLGISTAAAKKAIDIIDTASTIASIISLIGVVTGAGAISYAVVATAKAMIKKYGKKYAAAW |
| ACC_5 | *Clostridium* sp. | 81% | 64 | 6423 | 9.92 | 6 | 126.87 | 0.863 | LASTLGISSYAAKKVIDIIDAVSTVATIISLATAVIGAGAISAAIVATAKKMIKKYGKKYAAAW |
| ACC_6 | *Garciella nitratireducens* | 70% | 64 | 6508 | 9.90 | 5 | 131.09 | 0.970 | LASMLGISTVAAKKVIDIIDTASTVATIISLIGVIVGVGGITAGLVTTAKAMIKKYGKKYATMW |
| ACC_7 | *Sporolactobacillus vineae* | 67% | 64 | 6338 | 10.00 | 6 | 133.12 | 1.002 | LASTLGISTYAAKKAIDLISAASSIYAIIGLIAAIAGGGGIGIGILFTAKALIKKFGKKYAAAW |
| ACC_8 | *Marinitoga piezophila* | 64% | 64 | 6735 | 9.90 | 5 | 123.59 | 0.770 | LASMLGISKHAAEKVIDIIDTFSTVTTIISIITAVIGAGAITAGLVAVAKKMIKKYGKRYATMW |
| ACC_9 | *Helcococcus sp.* | 64% | 64 | 6415 | 10.00 | 5 | 132.81 | 1.014 | LTSTLGISAYAAKKIIDIISTASTIWSIVGIVAAVVGTGAIGVGILATAKTLVKKYGKAWAAAW |
| ACC_10 | *E. faecium, E. thailandicus* | 61% | 64 | 6317 | 9.90 | 5 | 134.38 | 0.953 | LTANLGISSYAAKKVIDIINTGSAVATIIALVTAVVGGGLITAGIVATAKSLIKKYGAKYAAAW |
| ACC_11 | *Kurthia sibirica* | 61% | 64 | 6649 | 9.70 | 4 | 134.06 | 0.991 | LAANLGISTAVATKIITIIDTYSTVTTIISLVGLLVGYGVFTTALVATAKTVISKYGKKYAVTW |
| ACC_12 | *Chloroflexota bacterium* | 58% | 64 | 6642 | 10.55 | 5 | 146.56 | 1.111 | LAANLGITTAAATRAIQIIDTASNIALIVSLLAVIFGAGVITTGLVITAKYVIRRFGKKYAIAW |

[a] Percent Identity relative to the reference variant (_1) of each bacteriocin group (query coverage varies among sequences).

[b] The grand average of hydropathicity (GRAVY) is a numerical index representing the overall hydrophobicity or hydrophilicity of a protein. GRAVY values > 0 indicate a predominantly hydrophobic protein, whereas GRAVY values < 0 indicate a predominantly hydrophilic protein.
